# ScoreNet: A Neural network-based post-processing model for identifying epileptic seizure onset and offset in EEGs

**DOI:** 10.1101/2020.12.21.423728

**Authors:** Poomipat Boonyakitanont, Apiwat Lek-uthai, Jitkomut Songsiri

## Abstract

We design an algorithm to automatically detect epileptic seizure onsets and offsets from scalp EEGs. The proposed scheme consists of two sequential steps: detecting seizure episodes from long EEG recordings, and determining seizure onsets and offsets of the detected episodes. We introduce a neural network-based model called *ScoreNet* to carry out the second step by better predicting the seizure probability of pre-detected seizure epochs to determine seizure onsets and offsets. A cost function called *log-dice loss* with a similar meaning to the F_1_ score is proposed to handle the natural data imbalance inherent in EEG signals signifying seizure events. ScoreNet is then verified on the CHB-MIT Scalp EEG database in combination with several classifiers including random forest, CNN, and logistic regression. As a result, ScoreNet improves seizure detection performance over lone epoch-based seizure classification methods; F_1_ scores increase significantly from 16–37% to 53–70%, and false positive rates per hour decrease from 0.53–5.24 to 0.05–0.61. This method provides clinically acceptable latencies of detecting seizure onset and offset of less than 10 seconds. In addition, an *effective latency index* is proposed as a metric for detection latency whose scoring considers undetected events to provide better insight into onset and offset detection than conventional time-based metrics.

## 1. Introduction

An epileptic seizure can be defined as a transient event of abnormal electrical activity in the brain (Fisher et al., 2014). Recently, epilepsy has affected approximately 65 million people around the world (Thurman et al., 2011). In clinical setting, neurologists can identify seizure characteristics through reviews of long scalp EEGs. However, the process of EEG visual examination is time-consuming, and prone to inconsistencies due to human errors brought by fatigue. To remove human errors from EEG-based seizure detection, we develop an automated epileptic seizure detection system that can label the time of seizure events in EEG signals. Several studies have developed methods to automatically detect epileptic seizures in EEG epochs (Acharya et al., 2015; Alotaiby et al., 2015; Boonyakitanont et al., 2020a; Hassan et al., 2020). Some studies focused on extracting single features relevant to EEG characteristics, *e*.*g*., amplitude (Satirasethawong et al., 2015; Shoeb and Guttag, 2010; Altunay et al., 2010), statistics (Samiee et al., 2015; Li et al., 2017), entropy (Tawfik et al., 2016; Li et al., 2018; Gupta and Pachori, 2019) and predictability (Kumar et al., 2015; Jaiswal and Banka, 2017; Li et al., 2019). Others have examined combinations of features to jointly distinguish between ictal patterns and normal activities (Mursalin et al., 2017; Alickovic et al., 2018; Vidyaratne and Iftekharuddin, 2017; Fergus et al., 2016). In addition, recent studies have favored deep learning models to examine EEG signals because these models can implicitly extract latent features and classify seizure episodes themselves (San-Segundo et al., 2019; Shoeibi et al., 2020; Takahashi et al., 2020; Shoeibi et al., 2021). Some studies have focused on the designs and choices of deep learning architectures suitable for indicating seizures (Acharya et al., 2018; Tian et al., 2019; Hu et al., 2020; OShea et al., 2020), while others have attempted to transform EEG segments into deep learning model inputs: *e*.*g*., an EEG plot image for a VGG16-based CNN model (Emami et al., 2019) and a three-dimensional EEG tensor for a temporal graph CNN model (Covert et al., 2019). Still, despite the promising performances achieved by these epoch-based seizure detection methods, they have not been able to realistically infer seizure onsets and offsets. Both false positive and false negative outcomes are possible in epoch-based methods, despite a lack of abrupt ictal pattern changes. An issue of this is exemplified in Figure 1, where false negatives lead to mislabeling of seizure onsets/offsets that incorrectly interpret one seizure episode as multiple events. Conversely, many isolated false positives will cause frequent false alarms. Therefore, it is still clinically inappropriate to determine seizure onsets and offsets as the first and last epochs of seizures by existing epoch-based seizure classifiers.

**Figure 1:**
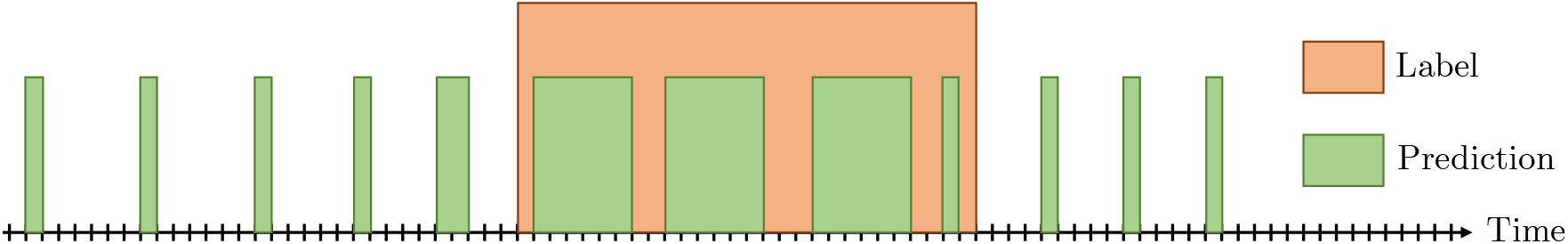
Misinterpretation of seizure events from isolated false positives and false negatives.

While seizure detection has seen wide interest, only a few researchers have focused on seizure onset and offset detection. Shoeb et al. (2011) proposed a method of detecting the endings of seizure episodes with a linear kernel support vector machine (linear SVM) using energies extracted from specific frequency bands of EEG epochs. However, this method failed to accurately determine seizure offsets when changes in seizure activity were gradual, and it needed a powerful seizure onset detector to satisfy its requirement of early seizure onset identification. Orosco et al. (2016) applied an artificial neural network (ANN) and linear discriminant analysis (LDA) with energy-based features calculated from stationary wavelet transform to determine seizures in EEG segments. Seizure onsets and offsets were then identified by the first and last indices of positive predictions during actual seizures. With discriminative features selected by a feature selection method called Lambda of Wilks, LDA outperformed ANN in the seizure detection. Nevertheless, no evidence could be found from the mean onset and offset latencies to indicate that the proposed method determines seizure onsets/offsets accurately, thus it is possible that the seizure events were correctly detected, but only for part of the duration of the actual event. Recently, a convolutional neural network (CNN) was proposed to spatially and temporally capture ictal patterns in EEG epochs (Boonyakitanont et al., 2020b). This method included an additional post-processing procedure designed from clinical characteristics of seizures to reduce the occurrence of false positives and false negatives from prior classification. As a result, this method had a significantly improved F_1_ score and drastically reduced FPR/h in almost all cases. However, the model parameters of this method were tediously manually selected for each patient which is a drawback that a self-learning scheme can address.

These issues encountered by previous epileptic seizure onset-offset detection methods were considered in the design of our automatic detection scheme which uses multi-channel scalp EEG signal inputs. Our detection system is divided into two sub-tasks: epoch-based classification, and onset-offset detection. The epoch-based classifier identifies seizures in small EEG segments independently and returns a predictive seizure probability as an output to the onset-offset detector. Several existing models including logistic regression, SVM, and decision tree can be applied in this first stage. Then, the second stage onset-offset detector receives as the input the preliminary results of epochs suspected of containing ongoing seizure occurrences, and estimates the starting and ending points of the detected events. This forms the first major contribution of our study: a novel model named *ScoreNet* that detects seizure onset-offsets by extending the detector structure in Boonyakitanont et al. (2020b). ScoreNet is a neural network-based model that automatically determines a group of seizure candidates from inputs, and then assigns a score to each candidate to determine the possibility that the candidate group should be regarded as a whole seizure activity. Such computations do not consider only a seizure epoch of interest, but also aggregate information from nearby epochs to take temporal changes of seizures into consideration. This feature makes ScoreNet unique from existing methods and improves the seizure detection performance from the first classification step. Moreover, since EEG seizure data are naturally highly imbalanced, the outcomes of existing detection methods tend to be biased towards a normal class. We address this issue by establishing a cost function in ScoreNet called *log-dice loss* based on a dice similarity coefficient.

Another issue in the abovementioned studies (Orosco et al. (2016); Chandel et al. (2019)) was that the mean latencies used for interpreting seizure onset-offsets were misleading; positive and negative latencies were defined as early and late predictions of seizure onset-offsets, respectively, and thus they could cancel each other out during the calculation of mean latencies. We address this issue by introducing the *effective latency index (EL-index)*, a novel time-delay metric that takes an undetected seizure onset-offset into consideration. Empirical results from this study will demonstrate that ScoreNet can dramatically reduce false positives and false negatives and precisely indicate seizure onsets and offsets from long EEG recordings. Additionally, it will be shown that using the log-dice loss helps overcome the class imbalance problem, and that the EL-index better represents the detection performance of a method than other conventional latency indices.

In summary, the contributions of this study are:

1. A neural network-based seizure onset and offset detector called *ScoreNet* and a loss function named *log-dice loss*.
2. A metric of onset-offset time delay termed the *effective latency index*.

This article is organized as follows: Section 2 presents the process of seizure onset-offset detection; Section 2.2 provides an in-depth explanation of ScoreNet; Section 2.3 provides the problem formulation including the proposed loss function; Section 3 describes the EL-index; Section 4 outlines all the experiments conducted to verify ScoreNet; finally, Section 5 presents the seizure classification and seizure onset-offset determination results, accompanied by discussions and with graphical illustrations.

## 2. Methods

### 2.1. Detection scheme

This research aspires to provide a method of detecting seizure episodes in long scalp EEGs and determining the onsets and offsets of the seizures. The detection process is divided into two sequential steps: epoch-based seizure classification, and seizure onset-offset detection, as shown in Figure 2a. Firstly, long multi-channel EEGs are segmented into small multi-channel EEG epochs to be used in seizure classification. A classifier receives raw data or a feature vector from epoch *i* as the input *x*_*i*_ and produces a seizure probability *z*_*i*_. An onset-offset detector then converts a collection of the prediction sequence *z* = (*z*_1_, *z*_2_, …, *z*_*N*_) into the modified sequence of seizure probabilities *ŷ* = (*ŷ*_1_, *ŷ*_2_, …, *ŷ*_*N*_) as a screened prediction of seizures. Finally, the onset and offset are indicated by the first and last indices of each predicted seizure that *ŷ*_*i*_ ≥0.5, respectively. Onset-offset detection is incorporated into the procedure to improve seizure detection accuracy upon the first seizure classification step.

**Figure 2:**
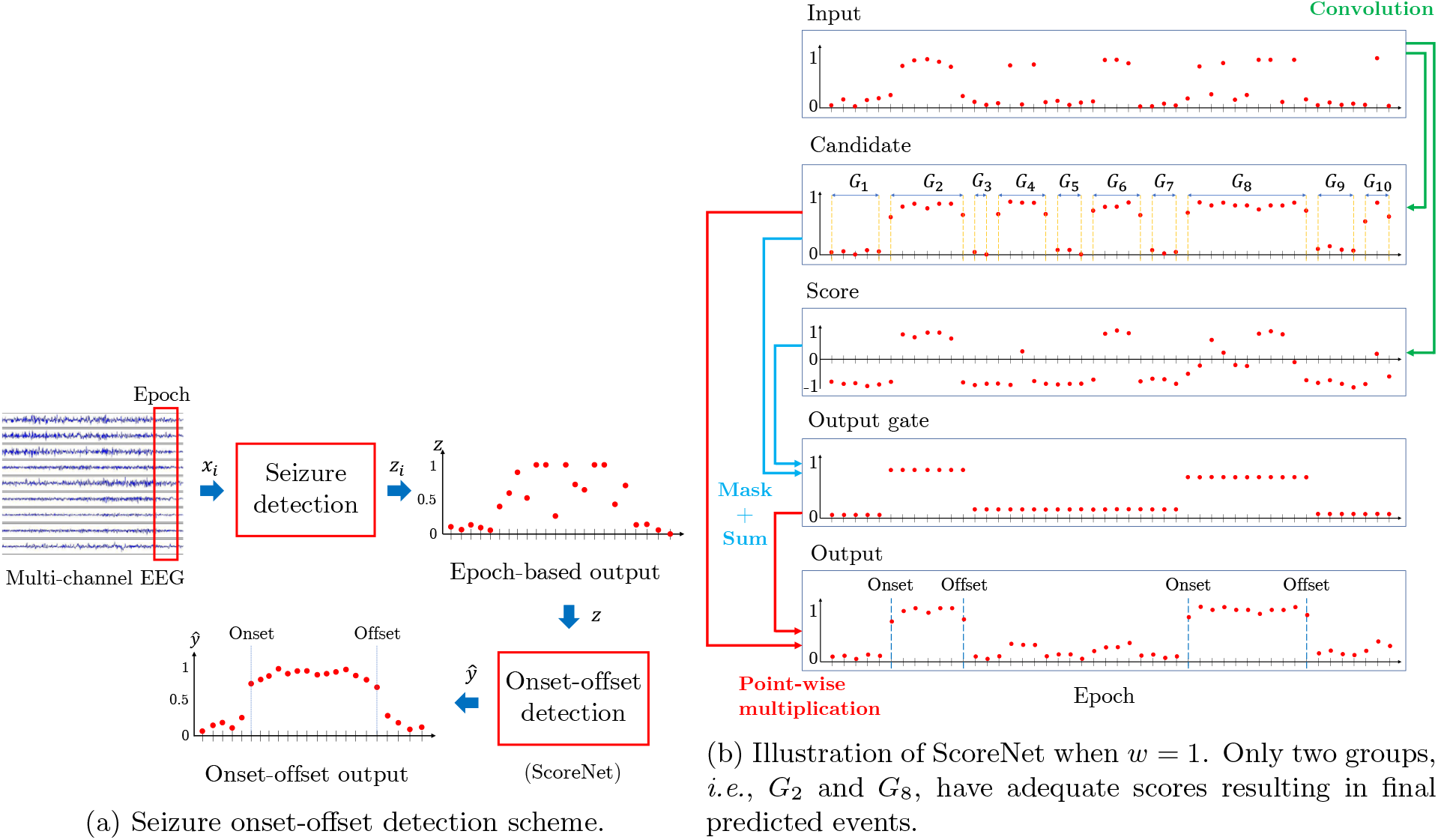
A scheme for determining the seizure onset and offset in long EEG signal. In this work, the onset and offset detection using ScoreNet is mainly focused.

### 2.2. ScoreNet

ScoreNet is a proposed onset-offset detector that takes epochs with detected likelihood of seizure as an input (*z*) and returns a screened detection of seizures in terms of probabilities as the output (*ŷ*), as shown in Figure 2a. A key to ScoreNet’s success is to use a convolution to connect adjacent epoch-based classification results into consideration at the same time. Moreover, the concept of automatically selecting a cell to be an output is used to predict a seizure event. Figure 2b shows ScoreNet’s computation process.

First, a *seizure candidate c*_*i*_ is defined as an epoch that has potentiality of containing a seizure by taking a convolution of neighbor epochs *z*_*i*_ with a linear filter *a*_1_ of length 2*w* + 1 and passing through the sigmoid function to return a value ranging from zero to one.

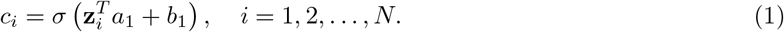

In the above, *σ* is a sigmoid function, **z**_*i*_ = (*z*_*i*+*w*_, …, *z*_*i*_, …, *z*_*i*−*w*_), and *z*_*j*_ = 0 for *j* ≤ 0 and *j > N*. Thresholding *c*_*i*_ gives a binary sequence, where values of *c*_*i*_ ≥*γ* are given a value of 1, and zero otherwise. The same consecutive values of the thresholded seizure candidate sequence are then clustered into groups, each denoted by *G*_*l*_. We also assign a degree to which each epoch influences the positive predictions of its neighbor epochs through a score *s*_*i*_, defined by

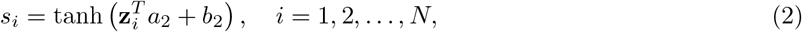

where *a*_2_ is a vector of length 2*w* + 1 and *b*_2_ is a bias term. Due to the hyperbolic tangent’s range, and through optimal selection of (*a*_2_, *b*_2_), we intend for *s*_*i*_ *<* 0 when **z**_*i*_ and its neighboring epochs are zero and *s*_*i*_ *>* 0 otherwise; thus, the score values can help distinguish between normal and seizure epochs. Then, each group of seizure candidates identified earlier are fed through a defined *output gate*, which is a nonlinear function of the averaged scores within a group to indicate the probability that the whole group is a seizure event, as in the expression

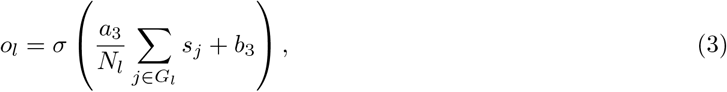

where *N*_*l*_ is the group size, *l* is the group index, *a*_3_ and *b*_3_ are scalar parameters.

Finally, ScoreNet produces an output *ŷ*_*i*_ which is a probability of seizure occurrence during seizure candidate *c*_*i*_, obtained by masking the seizure candidate with the output gate of group *G*_*l*_ to which *ŷ*_*I*_ belongs, as defined by

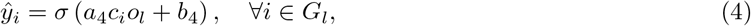

where *a*_4_ and *b*_4_ are scalar parameters. Note that the expressions of *c*_*i*_ and *s*_*i*_ resemble node equations in neural networks that take a linear combination of inputs and pass the combination through a nonlinear activation. The output gate equation (3) is also similar to an output gate in LSTM. For these reasons, the ScoreNet can be intuitively regarded as a neural network-based model.

As we have seen from the derivation of *ŷ*_*i*_ in (1)-(4), ScoreNet takes into consideration known temporal characteristics of seizures into its prediction of sequence *z* (series of epochs with suspected seizure activities) by convoluting *z* with linear filters (*a*_1_ and *a*_2_) and by using the knowledge of pre-determined seizure groups. By choosing optimal parameters, (*a*_*i*_, *b*_*i*_) for *i* = 1, …, 4, ScoreNet’s unprecedented approach of gathering information from neighboring EEG epochs should help it rule out unrealistic prediction outcomes, such as abrupt seizure activity changes within short durations.

Lastly, we note that ScoreNet is a more general framework extended from the counting-based approach in Boonyakitanont et al. (2020b). The model parameters were not optimally chosen in Boonyakitanont et al. (2020b), but instead set as *a*_1_ = *a*_2_ = **1**, *a*_3_ = *a*_4_ = 1, *b*_1_ = −1, *b*_2_ = −2, and *b*_4_ = 0, where **1** indicates the vector of ones with a compatible size, and only *b*_3_ can be tuned. Because Boonyakitanont et al. (2020b) used the Heaviside step function in (1), and with *z*_*i*_ being either zero (normal brain activity) or one (seizure), seizure candicate (*c*_*i*_) was obtained by counting seizures from adjacent epochs using a fixed threshold of *b*_1_. Similarly, Boonyakitanont et al. (2020b)’s treatment of (2), where neighboring epochs were counted, made it so that the score (*s*_*i*_) was set to one depending on whether the neighboring count exceeded a fixed threshold *b*_2_. Therefore, ScoreNet is expected to be an improvement over the counting-based method since its parameters can be optimized and selected.

### 2.3. Log-dice loss

To estimate parameters in ScoreNet, we will formulate a minimization of a loss function ℒ defined as the discrepancy between *y* – a binary sequence of annotated seizures (or target), and *ŷ* – a corresponding sequence of seizure probability generated by ScoreNet:

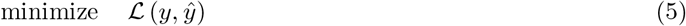

over the ScoreNet parameters (*a*_*i*_, *b*_*i*_) for *i* = 1, …, 4.

In a binary classification problem, many loss functions represent the similarity between *ŷ*_*i*_ and *y*_*i*_ in different forms. It is important to note that EEG seizure detection is an imbalanced classification because most EEG epochs are normal (*i*.*e*., *y*_*i*_’s are 0). For example, a cross-entropy loss is commonly employed for training a classifier in many classification problems, but it is not suitable for imbalanced classification (Lin et al., 2017) since it penalizes losses in both data classes equally. Instead, imbalanced data problems are handled by data manipulation (Nekooeimehr and Lai-Yuen, 2016; Ha and Lee, 2016; Jian et al., 2016; Haixiang et al., 2017), or by introducing a loss function that penalizes each data class differently. Some examples of uneven penalization are the weighting entropy (Krawczyk et al., 2014), or dice-coefficient-related loss (Lguensat et al., 2018; Milletari et al., 2016).

The dice similarity coefficient (DSC), or equivalently, the F_1_ score is a measure of the similarity between a predicted outcome and the ground truth, defined by

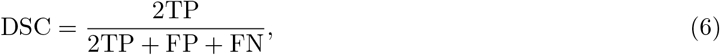

where TP, FN, and FP are the numbers of true positives, false negatives, and false positives, respectively. Notice that the DSC does not take the true negative (TN) into account. This implies that when the DSC is used in model training, and when the negative refers to the normal class, the model parameters are optimized to favor improving the accuracy of detecting positives (in our case: seizure events) over the majority class (normal brain activity). Other variants of DSC also exist and are used in imbalanced classification, such as the soft-dice loss by Lguensat et al. (2018) and the squared-dice loss^2^ by Milletari et al. (2016).

In this work, we establish the *log-dice loss* variant of DSC to tackle our imbalanced data problem, defined as

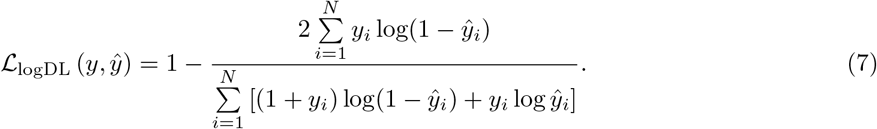

The log-dice loss is equal to 1 − DSC when substituting TP, FN, and FP in (6) for − Σ_*i*_ *y*_*i*_ log(1 −*ŷ*_*i*_), Σ_*i*_ *y*_*i*_ log *ŷ*_*i*_, and − Σ_*i*_(1 − *y*_*i*_) log(1 − *ŷ*_*i*_), respectively. Figure 3a shows the modified classification indices; by incorporating the logarithmic function, the index value rapidly increases where *ŷ* is close to *y*.

**Figure 3:**
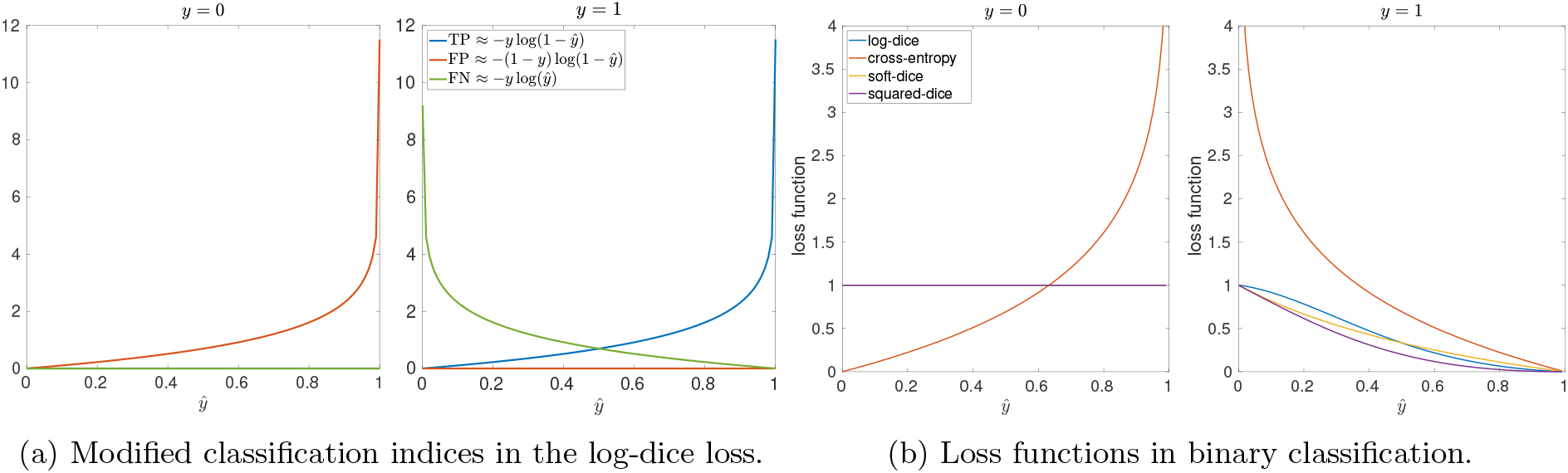
Comparisons of the log-dice loss with the others.

The value of ℒ_logDL_ is in the range of (0, 1] and decreases as *y* and *ŷ* become more similar. ℒ_logDL_ reaches its maximum of one (worst score) under two cases: i) when *y* = 0 (all samples are normal), regardless of the prediction *ŷ* because the index does not consider TN; or ii) when *y* = 1 and *ŷ* = 0 (no TP in the prediction). Figure 3b compares the cross-entropy, soft-dice, squared-dice and log-dice loss functions as *ŷ* varies under two values of *y* (one-sample case for illustration). When *y* = 0, ℒ_logDL_’s constant loss means that the normal class are neglected when optimizing model parameters. On the other hand, when *y* = 1, and *ŷ* ≤ 0.5, the log-dice loss has a higher penalty than the cross-entropy, soft-dice and squared-dice losses, implying that ℒ_logDL_ optimizes model parameters to prevent FN better than the other losses.

To solve (5), we apply a nonlinear conjugate gradient method because it has been shown to converge faster than other methods when training neural networks with limited variables (eight) to optimize (Le et al., 2011); the algorithm is described in Appendix A.

## 3. Effective Latency index

We propose the *effective latency index (EL-index)* as an indicator of delays between detected and actual onset-offsets, while also taking undetected events into account. The EL-index gives a zero (worst) score to any undetected event, and a positive score to any correctly detected event where the score increases as the delay decreases. Suppose there are *n* actual seizure activities and *k*_*i*_ is an indicator of event *i* being detected, *i*.*e*., *k*_*i*_ = 1 when the event is correctly detected, and *k*_*i*_ = 0 otherwise We also denote *d*_*i*_ *>* 0 and *d*_*i*_ *<* 0 for early and late detection of seizure onset and offset, respectively; note that *d*_*i*_ is not defined for an undetected seizure event. For a given 0 *< r <* 1, the EL-index is defined as

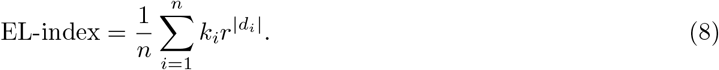

Index values can range from zero (missing all seizure events) to one (perfectly detecting all events). A large latency in the detection of any event will cause an exponential decrease in the EL-index at a decay rate of *r*. If we denote GDR (good detection rate) as a portion of correctly detected seizure events in one record given by 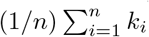, then the EL-index can be regarded as an exponentially weighted GDR, and that it satisfies the bounds:

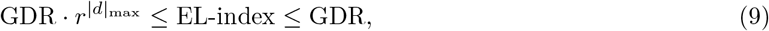

where |*d* | _max_ is the maximum value of absolute time delays. It is evident that the EL-index cannot be higher than GDR, while also being lower-bounded by the function GDR *r*^|*d*|^. Moreover, a relation between the EL-index and the mean absolute latency (MAL) can be derived and provides some connection about empirical distributions of collected time delays. Suppose we have time delay samples from two test results that have the same MAL and GDR; however, the first set contains narrowly distributed time delays, while the samples of the second set are highly varied. For the first set, when all time delays are similar, *i*.*e*., *d*_*i*_ ≈ *±*MAL, for any detected event *i*, the EL-index can be approximated by

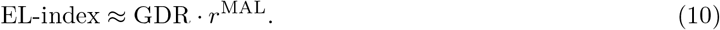

In contrast, when the time delays are widely spread, the approximation (10) does not hold, and the EL-index is always higher than in cases of narrowly distributed latencies:

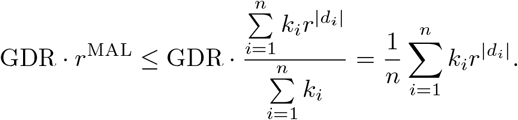

We will use a hypothetical example illustrated in Figure 4 to demonstrate the EL-index’s superiority in describing latency error in detected signals over using the mean latency as in Orosco et al. (2016); Chandel et al. (2019). The values in Figure 4 shows the comparison between the mean latency, MAL, and EL-index when *r* = 0.9. While methods A and B both have zero mean latencies of onset detection in this hypothetical example, neither perfectly detected seizure onsets; hence, this example shows that the mean latency can give a false sense of accuracy in the detection method. Next, if MAL is considered instead of the mean latency, method A still suggests higher detection accuracy (less latency and smaller MAL value) than method B, even though method A completely missed detecting the second seizure event (*k*_2_ = 0). Once again, this example has shown that even MAL is not a good indicator of detector latency if there are many undetected events. However, the EL-index in this example does reveal that method B performs better than method A in detecting onsets, because the EL-index considers the fact that method A did not detect the second seizure event. Also, looking at Figure 4, it is apparent that method A detects onsets more accurately than offsets, which is correctly indicated by method A’s higher EL-index of detecting onsets than offsets.

**Figure 4:**
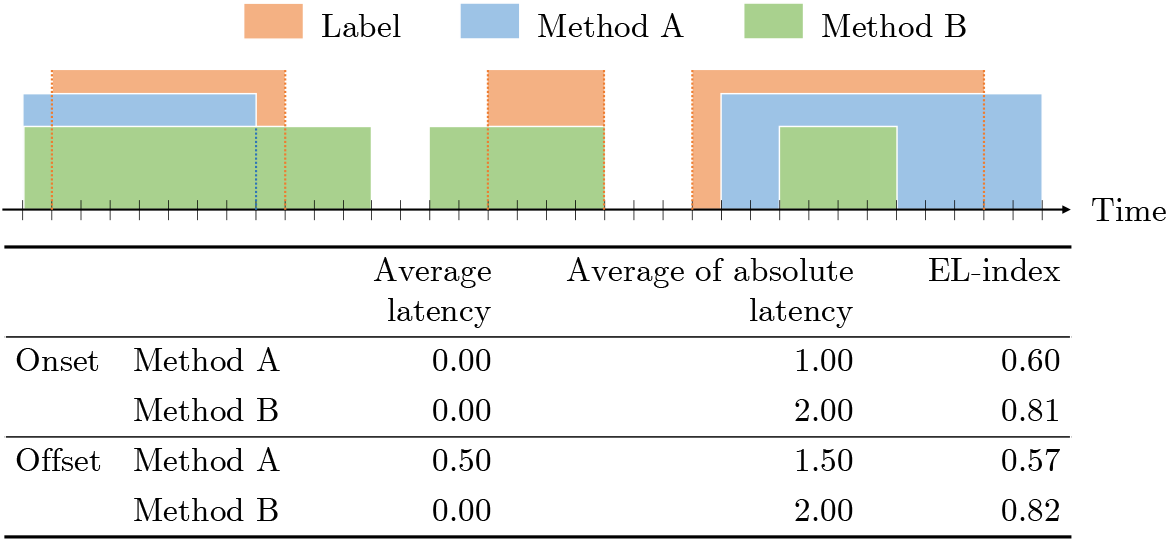
Example of time-based measurements. Orange segments are annotated EEG epochs, while blue and green segments are predicted seizures from method A and B, respectively.

## 4. Experiment

### 4.1. Data preparation

The CHB-MIT Scalp EEG database is a collection of EEG signals observed from 24 pediatric cases at the Childrens Hospital Boston (Goldberger et al., 2000; Shoeb, 2009); it is available on PhysioNet (https://physionet.org/content/chbmit/1.0.0/). The data were sampled at a rate of 256 Hz and a resolution of 16 bits; they were recorded with the international 10–20 electrode system. Most records consist of 23 channels collected from either referential or bipolar montages; for more details of the number of records, please see Table C.4 in Appendix C. Since the data were recorded with both referential and bipolar montages, the EEG records were firstly rearranged to the bipolar montage system. The sequential order of the modified 18 channels were *FP1-F7, F7-T7, T7-P7, P7-O1, FP1-F3, F3-T3, T3-P3, P3-O1, FP2-F4, F4-C4, C4-P4, P4-O2, FP2-F8, F8-T8, T8-P8, P8-O2, FZ-CZ*, and *CZ-PZ*. The long EEG records were segmented into non-overlapping epochs of one second.

### 4.2. Evaluation

Seizure detection methods were evaluated using a patient-specific leave-one-record-out cross validation (LOOCV) scheme, where *all records* in the CHB-MIT Scalp EEG database were used. Training and test data were taken from the same patient but different records. Detection methods were validated with all patient cases; they were assessed using event-based, epoch-based, and time-based metrics (Temko et al., 2011; Boonyakitanont et al., 2020a). The event-based metrics included good detection rate (GDR) and false positive rate per hour (FPR/h). The epoch-based metric was F_1_, which fairly assesses seizure detection results as it is an imbalanced classification (Powers, 2011). The time-based metric was the EL-index as explained in Section 3. We also used our detection results to compare the EL-index with other latency indices. If one true event was interpreted as more than one event by a detector, then the onset of the *first* predicted event and the offset of the *last* positive activity were used to calculate the time-based metrics.

### 4.3. Experimental Setup

The first-stage seizure detection in Figure 2a was performed by five classifiers: CNN, logistic regression, linear SVM, decision tree, and random forest. For CNN, we adopted the structure in Boonyakitanont et al. (2020b) that can extract meaningful features from raw EEG segments that it uses as the input. The other classifiers depended on widely-used features that characterize ictal and normal patterns as reported in Boonyakitanont et al. (2020a); Shoeb and Guttag (2010); Acharya et al. (2015, 2013). *Time-domain features* computed from raw EEG epochs were the variance, energy, nonlinear energy, Shannon entropy, sample entropy, and approximate entropy. *Frequency-domain features* calculated from power spectral densities were the energies from eight sub-bands in the range of 0–25 Hz. In addition, *time-frequency-domain features* were extracted with discrete wavelet transform coefficients from five decomposition levels with the Daubechies 4 tap wavelet; these features were the mean absolute value, variance, energy, maximum, minimum, and line length. Features were extracted from each channel in an EEG epoch and normalized to a *z*-score. In total, 900 features were combined into a feature vector for the classification of EEG epochs.

As for the second-stage onset-offset detection of Figure 2a, ScoreNet and the counting-based method of Boonyakitanont et al. (2020b) were used and compared, where the sizes of *a*_1_ and *a*_2_ in both implementations were set to 13. ScoreNet was trained with binary cross-entropy, soft-dice loss (Lguensat et al., 2018), and square-dice loss (Milletari et al., 2016) for comparisons of dealing with an imbalanced data problem. Codes and results are available at https://github.com/Siyaosk129/ScoreNet.

## 5. Results and Discussions

Figures 5a and 5b show the GDR and FPR/h scores of every seizure-detection method we tested. Recall that our seizure-detection scheme was divided into two steps. The first-stage epoch-based classification was performed by CNN, logistic regression, linear SVM, decision tree, and random forest classifiers; GDR and FPR/h scores of performing only the first-stage classification with each of these classifiers are labeled as `classification`. The second step involved applying ScoreNet with different loss functions on the first-stage results; the loss functions used are labeled as in the parentheses as follows: entropy (`entropy`), soft-dice (`softdl`), squared-dice (`sqdl`), and log-dice (`logdl`). When the counting-based method was applied, we labeled these results as `counting`. We will describe and compare the results in three folds: the improvement of using onset-offset detection over epoch-based classification, a comparison between the counting-based and ScoreNet methods, and the accuracy of the onset-offset determination.

**Figure 5:**
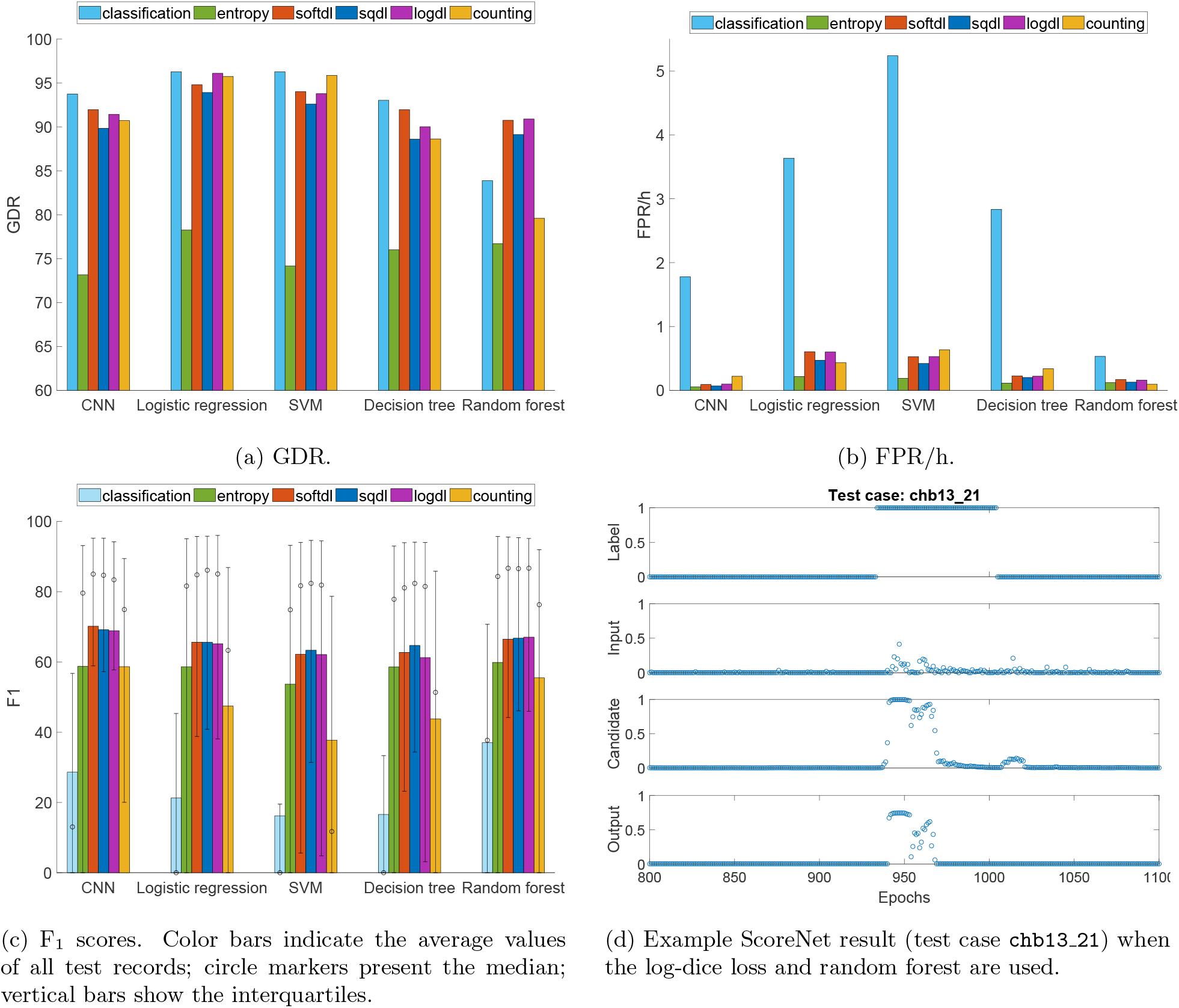
Comparisons of classification indices obtained from test cases using different epoch-based seizure detection methods.

### 5.1. Seizure detection

We can see the improved seizure detection results of applying ScoreNet over using only the classifiers in terms of F_1_, GDR and FPR/h scores in Figures 5a to 5c. Alone, the seizure classifiers detected seizure events at GDRs of more than 80%, but obtained F_1_ of less than 40%, and had drastically varying FPR/h between 0.53 to 5.24 on average. If we consider all these scores as overall performance metrics, then random forest and CNN are both compromised classifiers that are good at detecting most seizure events, but at a cost of frequent erroneous inferences; both of these classifiers have room for improvement.

By applying ScoreNet with any of the loss functions, regardless of the classification methods, F_1_ increased at least 18% over lone classifier F_1_ scores, and FPR/h significantly reduced at least 0.36 times per hour; however, GDRs did drop slightly, except in the case of random forest where the GDR increased up to 7%. The favorable outcomes afforded by combining random forest and ScoreNet are exemplified by the example case illustrated in Figure 5d. Random forest produced an interval of small seizure probabilities that were sufficiently distinguishable from the background for ScoreNet to recognize them as seizure events and significantly boost the magnitudes of their candidate and output scores. Thus, this confirmed that using ScoreNet, with any dice loss function, generally improves epoch-based classification performances; specifically, ScoreNet can indicate some seizure events that are not detected by a lone classification process. Secondly, this also confirmed that dice loss functions are appropriate for handling imbalanced-class data problems.

Since the application of ScoreNet resulted in FPR/h and F_1_ score improvements but GDR drop-offs, we can generally interpret that FP, FN, and TP decreased with ScoreNet present, and the extent to which these predictions were eliminated depended on the employed cost function. For example, using ScoreNet with the cross-entropy loss, which mainly penalizes errors of normal/majority class samples, resulted in a large reduction of FPR/h. This was the result of ScoreNet with cross-entropy reducing several isolated FPs at the expense of also unfavorably removing some TPs, where perhaps the seizures only occurred within a few epochs that did not generate enough predicted positives for event detection.

For dice loss functions, ScoreNet generally yields similar seizure detection performances across any seizure detection methods. ScoreNet with the soft-dice loss provided the best general seizure-activity detection results, although ScoreNet with the squared-dice loss provided better (lower) FPR/h. In addition, using the log-dice loss improved classification performance when classification errors were large. Referring again to the results of using random forest and log-dice loss in Figure 5d, random forest was initially unable to detect the seizure in epochs 940–1000, where the seizure probability inputs were smaller than 0.5. However, by incorporating ScoreNet, the seizure predictions in epochs 940–1000 were boosted enough that these epochs ended up having active seizure labels as well. Thus, it was the log-dice loss’s ability to uncover previously undetected seizure activities that resulted in better GDR and F_1_ scores, and made it the form of dice-loss that most improved the random forest classifier.

Table 1 summarizes previous studies’ seizure detection performances, with the amount of data and validation schemes they used. In our study, the highest F_1_ score was achieved by ScoreNet with soft-dice loss and CNN as the prior classifier; therefore, we compared the performance of this method (`proposed method`) to the previous references. It was difficult to directly compare the performance metrics, since the validation schemes and data selection were different among these studies. Many of the previous studies selected specific records from the database, and only a few applied LOOCV as a validation scheme. Tang et al. (2020) used only records containing seizures in the experiment, and Yuan et al. (2017) did not report on their data specification nor validation scheme. Moreover, some studies performed data sampling to create balanced training data (Hassan et al., 2020), but in practice some EEG characteristics – such as rarity and types – cannot be accurately selected; it is more clinically challenging to use all data and apply LOOCV to verify the detection performance. With these considerations in mind, our proposed method yielded competitive performances against previous results.

**Table 1:** Comparison of seizure detection methods using CHB-MIT database.

| Reference | Detection type | Data specification | Validation | Method | Acc | Sen | Spec | F<sub>1</sub> | GDR | FPR/h |
| --- | --- | --- | --- | --- | --- | --- | --- | --- | --- | --- |
| Satirasehawong et al. (2015) | Event | Long seizures | No CV | aEEG + adaptive threshold | NR | NR | NR | NR | 88.50 | 0.18 |
| Yuan et al. (2017) | Event | NR | NR | Spectrogram+mSSDA | 93.82 | NR | NR | 96.05 | NR | NR |
| Vidyaratne and Iftikharuddin (2017) | Event | Specific records (22 cases) | LOOCV | Energy and fractal | NR | NR | NR | NR | 97.00 | 0.10 |
| Gao et al. (2020) | Event | 166 mins (11 cases) 70% training data | No CV | PSD + CNN-based | 92.60 | 92.30 | 97.00 | NR | NR | NR |
| Hu et al. (2020) | Event | All, Balanced training data | No CV | Statistical features + Bi-LSTM | 92.66 | 93.61 | 91.85 | NR | NR | NR |
| Shoeb and Guttag (2010)* | Onset | NR | LOOCV | Energy + RBF SVM | NR | NR | NR | NR | 96.00 | 0.08 |
| Tang et al. (2020) | Onset | Seizure in record | LOOCV | Unified multi-level spectral-temporal feature + RBF SVM | 97.80 | NR | NR | 78.00 | 97.20 | 0.64 |
| Orosco et al. (2016) | Onset/offset | Bipolar montage (18 cases), Balanced training data | 10-fold | Relative band energy+LDA | NR | NR | 99.99 | NR | 92.60 | 0.30 |
| Chandel et al. (2019) | Onset/offset | 397 hrs (18 cases), 60% training data | 5-fold | Statistical features+LDA | 98.00 | NR | 98.05 | NR | 100.00 | 4.02 |
| Boonyakitanont et al. (2020b) | Onset/offset | All | LOOCV | CNN+counting-based method | 99.72 | 72.78 | 99.82 | 64.40 | 83.41 | 0.12 |
| **Proposed method** | Onset/offset | All | LOOCV | CNN+ ScoreNet (soft-dice loss) | 99.83 | 76.54 | 99.92 | 70.15 | 91.96 | 0.09 |

NR = no report, All = use full data set, \* Use median instead of mean in the report

### 5.2. Onset and offset determination

The GDR, MAL, and corresponding EL-indices of every test results are displayed in Figure 6. Although delay is not defined when the GDR = 0%, we have set *d* to zero (indicated by yellow markers) in these cases for illustrative purposes. There were a significant number of test cases with 0% GDR, or undetected events. Hence, using only the MAL as a performance metric would mistakenly ignore these detection failures, whereas the EL-index captures undetected events since 0% GDR is still reflected by a zero EL-index score. For low GDR cases (about 40–50%), seizure events appeared to be randomly detected, which still resulted in misleadingly low MAL scores in some test samples, but were properly reflected by the decreased EL-index scores (indicating worse performance). These observations suggest that the proposed EL-index is more suitable as a time-based index than the MAL.

**Figure 6:**
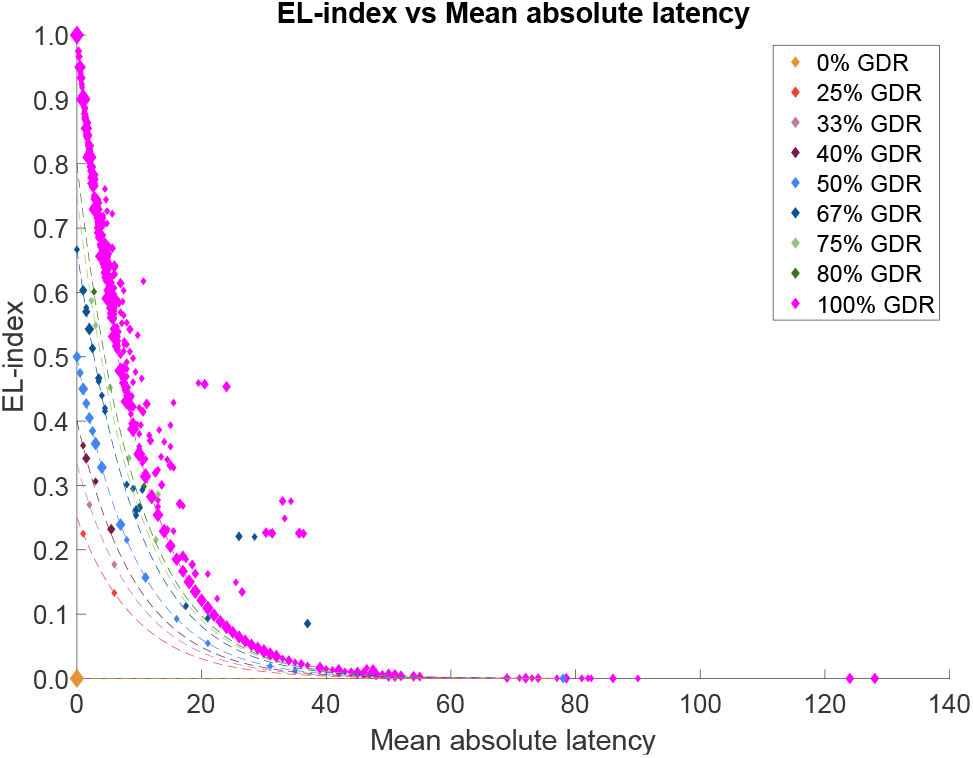
Relation of EL-index and mean absolute latency (MAL) from the test data given *r* = 0.9. Marker sizes are proportional to the number of samples in a log scale. Dashed line illustrate GDR *· r*^|*d*|^ limits.

For any non-zero GDR case, the EL-index was biased to have higher values when many seizure events are detected, and the latencies in those detected seizure onset/offsets are insignificant. In Figure 6, the relationship between EL-index and MAL mostly satisfies the exponential bound (9), GDR *· r*^|*d*|^ represented by dashed lines. As analyzed in Section 3, this means that latencies in detected seizure onset/offsets generally had low variation. In addition, the MAL cannot provide information on whether different cases with similar MAL scores have similar or different onset-offset latencies, but the EL-index values can. Therefore, the EL-index provides not only insight into the accuracy of seizure onset-offset detection, but also the accuracy of seizure event detection, and interpretation of the empirical distributions of latencies when considered jointly with the GDR.

Figure 7 compares the performances of detecting onsets versus offsets as measured by the EL-index with *r* = 0.9. In the case of seizure onset detection, the mean EL-indices were in the range of 0.50–0.71 and the medians ranged from 0.59 to 0.81. As for detecting seizure offset, the mean and median ranges of EL-indices were slightly lower 0.45–0.67 and 0.53–0.73, respectively. As the minimum median (among several classifiers) was around 0.53, we can interpret together with Figure 6 that around half of the test samples were detected seizures with a MAL of less than 10 seconds, which is clinically acceptable.

**Figure 7:**
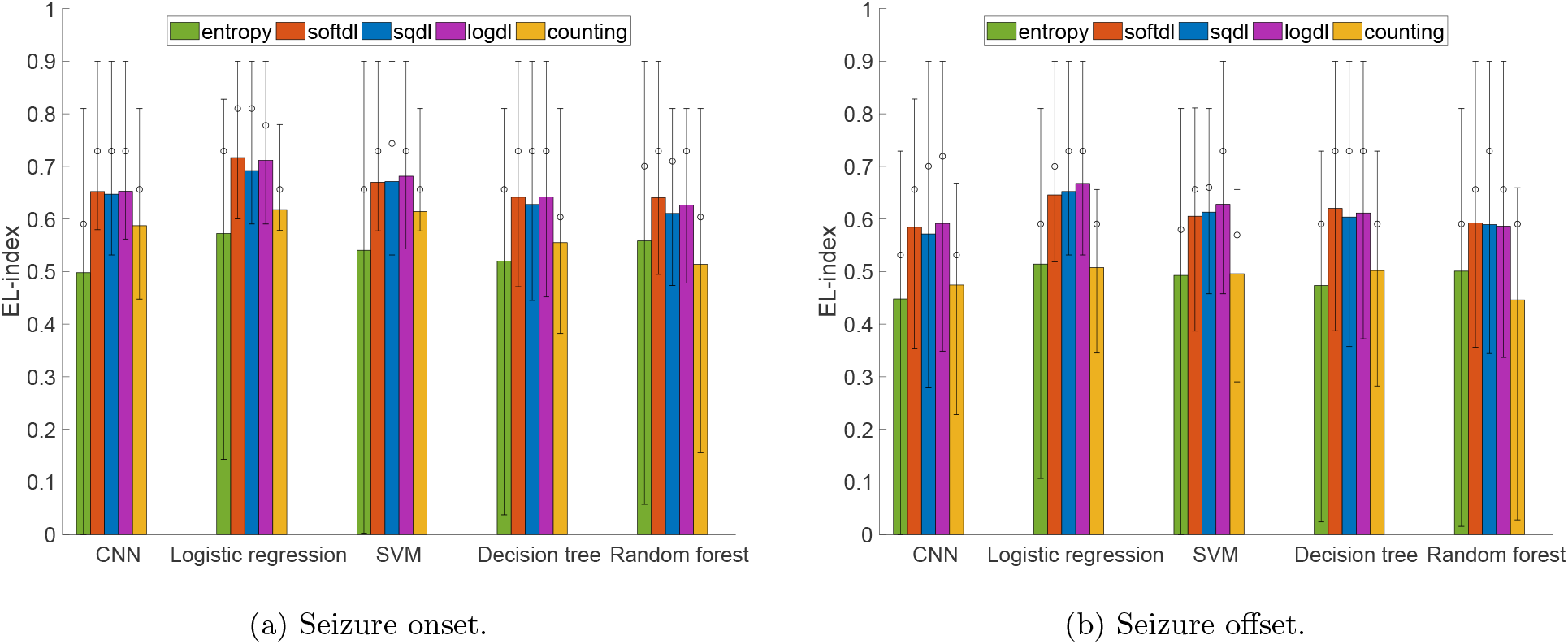
Onset-offset detection performances measured by the EL-index. The color bars indicate averaged values; circle markers present the median; vertical bars show the interquartiles.

Table 2 compiles the EL-indices obtained from the detection methods with various choices of loss functions and classifiers. The EL-indices obtained by using the dice loss functions were similarly high compared to those of the entropy loss and the counting-based method; in particular, the log-dice loss achieved slightly better EL-indices than other dice loss functions. Also, all methods indicated the seizure onsets better than the seizure offsets. This is due to the characteristics of epileptic seizures where ictal patterns occurring at the event ending are typically less dominant than the patterns at the beginning, and so it is harder for detectors to distinguish seizure epochs near the end of events. Recall that the log-dice loss can fix classification outcomes that are erroneously labelled as negatives due to low seizure probabilities; we now also know that these errors tend to occur at the end of seizure event, and we should see the most performance improvements when employing the log-dice loss in cases where large numbers of these classification errors occur.

**Table 2:** The average EL-index of seizure onset and offset determination. For each classifier, the boldface value is the maximum EL-index.

|  |  | CNN | Logistic<br>regression | SVM | Decision<br>tree | Random<br>forest |
| --- | --- | --- | --- | --- | --- | --- |
| Onset | entropy | 0.50 | 0.57 | 0.54 | 0.52 | 0.56 |
|  | softdl | **0.65** | **0.72** | 0.67 | **0.64** | **0.64** |
|  | sqdl | **0.65** | 0.69 | 0.67 | 0.63 | 0.61 |
|  | logdl | **0.65** | 0.71 | **0.68** | **0.64** | 0.63 |
|  | counting | 0.59 | 0.62 | 0.61 | 0.55 | 0.51 |
| Offset | entropy | 0.45 | 0.51 | 0.49 | 0.47 | 0.50 |
|  | softdl | 0.58 | 0.65 | 0.61 | **0.62** | **0.59** |
|  | sqdl | 0.57 | 0.65 | 0.61 | 0.60 | **0.59** |
|  | logdl | **0.59** | **0.67** | **0.63** | 0.61 | **0.59** |
|  | counting | 0.47 | 0.51 | 0.50 | 0.50 | 0.45 |

## 6. Conclusion

This research established an automatic epileptic seizure onset and offset detection scheme composed of two processes: detecting seizures in epochs from EEG signals, and determining the beginning and ending points of a seizure event. ScoreNet was designed to detect epileptic seizure onsets and offsets from the first-stage epoch-based classification results. It incorporates a log-dice loss function to handle the data class imbalance that is inheritent in using EEG signals to classify seizure events. Its ability to detect seizure onset/offsets was demonstrated by a proposed EL-index. The proposed scheme was evaluated with real patient cases from the CHB-MIT Scalp EEG database. In handling these cases, ScoreNet performed better than a lone epoch-based seizure classification method with improved F_1_ scores of up to 70.15%, dramatically reduced the false alarms rates to 0.05 times per hour, and yielded onset-offset detection errors of typically less than 10 seconds, which are clinically acceptable. Performance improvements yielded using the log-dice loss were most pronounced when prediction errors from epoch-based seizure detection were large. In addition, the EL-index was proven to be suitable for measuring seizure onset/offset detection latencies, as it provides information about both the correct detection of seizure events, and the latency distribution.

While our method provides several advantages over previous seizure detection methods, some limitations and concerns with it are worth noting. First, we employed a patient-specific scheme in this paper to prove ScoreNet’s ability to improve seizure detection performance. As such, the diversity of seizure types and artifact degrees in different patients may have been left out of consideration if ScoreNet’s enhancement ability is unique to each patient. Since ScoreNet has to be specifically trained for each patient, in order to be used in a clinical setting, EEG recordings and seizure annotations must first be collected from a patient to initially train the model. This limits the models practicality, raising the concept of a universally pre-trained detector. Perhaps ScoreNet can be trained across patients or on a larger dataset, but how the average detection performance would respond is yet to be seen, especially considering the high EEG variations between patients. Hence, perhaps future work can focus on developing a universal seizure detector that exploits common spatial and temporal patterns of seizures across various patients.

## 7. Acknowledgment

This work is financially supported by the 100th Anniversary Chulalongkorn University Fund for Doctoral Scholarship, the 90th Anniversary of Chulalongkorn University Fund (Ratchada-phiseksomphot Endowment Fund), and the 2020 Chula Engineering Research grant.

## Appendix A. Algorithm of ScoreNet

Here, we describe pseudo codes of the nonlinear conjugate gradient method that trains ScoreNet in (5). We note that a loss function ℒ(*y, ŷ*) is in fact a function of problem data (*y*) and the ScoreNet parameters; refer to the description of *ŷ* in (1)-(4). For optimization purpose, it is better to equivalently present the loss function as ℒ (*y, ŷ*) ≜ *f* (*x*) where *x* = (*a*_1_, *b*_1_, *a*_2_, *b*_2_, *a*_3_, *b*_3_, *a*_4_, *b*_4_) is the ScoreNet parameter vector. As required by the algorithm, computing the gradient of *f* can be facilitated by arranging ℒ (*y, ŷ*) as the sum of local losses:

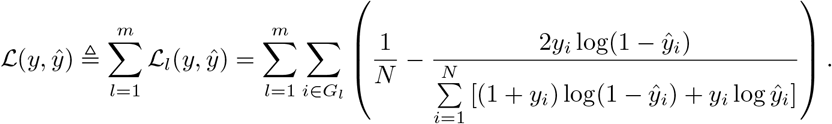

Hence, the gradient of *ℒ*_*l*_(*y, ŷ*) for *l* = 1, …, *m* can be computed in parallel. As for *∇ℒ*_*l*_, the derivatives with respect to *b*_4_ and *a*_4_ are carried out first using the chain rule. Similar to the backpropagation in neural networks,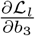 can be expressed as a function of 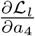, and 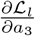 as a function of 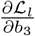. Applying the straightforward chain rule, we also obtain the remaining derivatives where 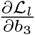 is repeatedly used so it can be cached once in the computation.

### Algorithm 1

Training ScoreNet

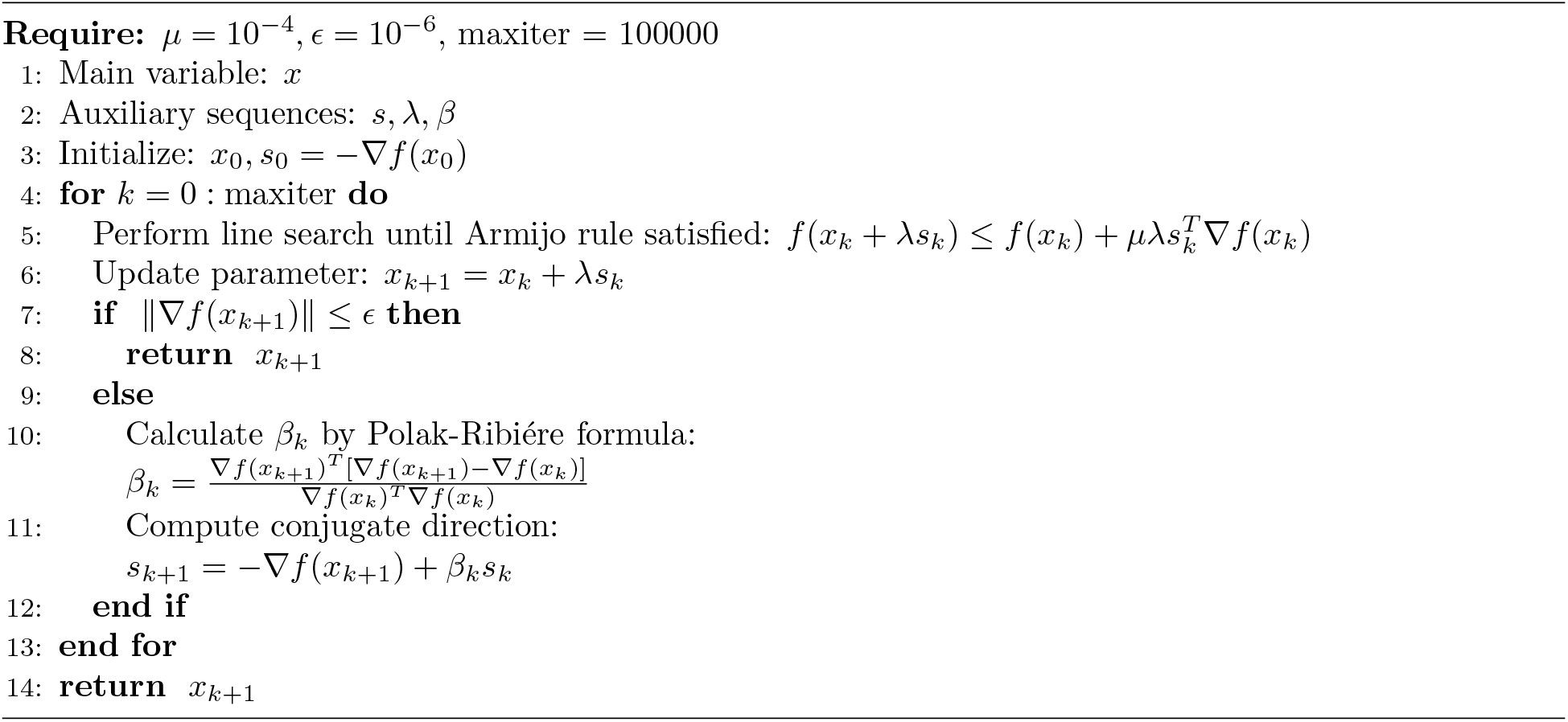

Algorithm 1 returns the ScoreNet parameters that (locally) minimizes the loss function. It was implemented for all choices of the loss function (cross-entropy loss, soft-dice loss, squared dice loss and log-dice loss) in this work.

## Appendix B. Initial ScoreNet parameters

Algorithm 1 requires an initial set of ScoreNet parameters as *x*_0_. An issue of optimizing the ScoreNet parameters is that since *z*_*i*_ is the output of a classification problem of highly imbalanced data, *z* is extremely sparse; **z**_*i*_ is usually a zero vector, and the gradients vanish. Thus, the parameters *a*_1_ and *a*_2_ only slightly adjust with consecutive iterations, and they hardly ever converge to a good local minimum. Therefore, we selected three sets of *x*_0_ according to prior classification results based on training data. We initialized *a*_1_ and *a*_2_ based on the three situations, and the initialization of the other parameters were empirically chosen to avoid landing in a poor local optimum. The selection of initial *a*_1_ and *a*_2_ follows the parameters of the counting based method, *i*.*e*., multiples of **1**. When there are many false negatives from the first-stage seizure-epoch classification step, the magnitudes of *a*_1_ and *a*_2_ will be high to boost a seizure candidate *c*_*i*_ and easily assign *s*_*i*_ = 1. On the other hand, when the number of isolated false positives is high, the magnitudes of *a*_1_ and *a*_2_ will be small to suppress the effect of false positives. In between, *a*_1_ and *a*_2_ should have intermediate magnitudes if the numbers of false positives and negatives are more similar. The choices of initial parameters are summarized in Table B.3.

**Table B.3:** Initial ScoreNet parameters for training. Vector of ones is represented by **1**.

| Cases | Parameters |  |  |  |  |  |  |  |
| --- | --- | --- | --- | --- | --- | --- | --- | --- |
| | $a_1$ | $a_2$ | $a_3$ | $a_4$ | $b_1$ | $b_2$ | $b_3$ | $b_4$ |
| High FP | $\mathbf{1}$ | $\mathbf{1}$ | 3 | 3 | -1 | -2 | 0 | -1 |
| High FN | $9 \cdot \mathbf{1}$ | $9 \cdot \mathbf{1}$ | 4 | 4 | -4 | -8 | 1 | -1 |
| Balance | $6 \cdot \mathbf{1}$ | $6 \cdot \mathbf{1}$ | 3 | 3 | -3 | -9 | 0 | -1 |

## Appendix C. Database summary

A record summary of the CHB-MIT Scalp EEG database is shown in Table C.4.

**Table C.4:** Summary of the CHB-MIT Scalp EEG database.

| Cases | # records | Total duration (sec) | # seizures | Seizure duration (sec) |
| --- | --- | --- | --- | --- |
| chb01 | 42 | 145,988 | 7 | 449 |
| chb02 | 36 | 126,959 | 3 | 175 |
| chb03 | 38 | 136,806 | 7 | 409 |
| chb04 | 42 | 561,834 | 4 | 382 |
| chb05 | 39 | 140,410 | 5 | 563 |
| chb06 | 18 | 240,246 | 10 | 163 |
| chb07 | 19 | 241,388 | 3 | 328 |
| chb08 | 20 | 72,023 | 5 | 924 |
| chb09 | 19 | 244,338 | 4 | 280 |
| chb10 | 25 | 180,084 | 7 | 454 |
| chb11 | 35 | 123,257 | 3 | 809 |
| chb12 | 24 | 85,300 | 40 | 1,515 |
| chb13 | 33 | 118,800 | 12 | 547 |
| chb14 | 26 | 93,600 | 8 | 177 |
| chb15 | 40 | 144,036 | 20 | 2,012 |
| chb16 | 19 | 68,400 | 10 | 94 |
| chb17 | 21 | 75,624 | 3 | 296 |
| chb18 | 36 | 128,285 | 6 | 323 |
| chb19 | 30 | 107,746 | 3 | 239 |
| chb20 | 29 | 99,366 | 8 | 302 |
| chb21 | 33 | 118,189 | 4 | 203 |
| chb22 | 31 | 111,611 | 3 | 207 |
| chb23 | 9 | 95,610 | 7 | 431 |
| chb24 | 22 | 76,640 | 16 | 527 |
| **sum** | **686** | **3,536,540** | **198** | **11,809** |

## Footnotes

2 The actual function name in Milletari et al. (2016) is the *soft-dice loss*. We refer to it by a different name to avoid confusion.

